# Prevention and therapy of SARS-CoV-2 and the B.1.351 variant in mice

**DOI:** 10.1101/2021.01.27.428478

**Authors:** David R. Martinez, Alexandra Schaefer, Sarah R. Leist, Dapeng Li, Kendra Gully, Boyd Yount, Joy Y. Feng, Elaine Bunyan, Danielle P. Porter, Tomas Cihlar, Stephanie A. Montgomery, Barton F. Haynes, Ralph S. Baric, Michel C. Nussenzweig, Timothy P. Sheahan

## Abstract

Improving the standard of clinical care for individuals infected with SARS-CoV-2 variants is a global health priority. Small molecule antivirals like remdesivir (RDV) and biologics such as human monoclonal antibodies (mAb) have demonstrated therapeutic efficacy against SARS-CoV-2, the causative agent of COVID-19. However, it is not known if combination RDV/mAb will improve outcomes over single agent therapies or whether antibody therapies will remain efficacious against variants. In kinetic studies in a mouse-adapted model of ancestral SARS-CoV-2 pathogenesis, we show that a combination of two mAbs in clinical trials, C144 and C135, have potent antiviral effects against even when initiated 48 hours after infection. The same antibody combination was also effective in prevention and therapy against the B.1.351 variant of concern (VOC). Combining RDV and antibodies provided a modest improvement in outcomes compared to single agents. These data support the continued use of RDV to treat SARS-CoV-2 infections and support the continued clinical development of the C144 and C135 antibody combination to treat patients infected with SARS-CoV-2 variants.

## INTRODUCTION

A novel human coronavirus, SARS-CoV-2, emerged in late 2019 in Wuhan, China (Zhou et al., 2020b; Zhu et al., 2020) as the causative agent of coronavirus disease 2019 (COVID-19). The spread of SARS-CoV-2 was explosive with ~140 million confirmed cases and >3 million deaths worldwide as of April 2021. Few therapies are available to treat COVID-19 disease in humans and the rapid evolution of SARS-CoV-2 variants threatens to diminish their efficacy. Remdesivir (RDV, Veklury) is the only U.S. Food and Drug Administration (FDA) approved direct-acting, small molecule antiviral to treat COVID-19. Prior to the emergence of SARS-CoV-2, RDV showed broad-spectrum activity against highly pathogenic human coronaviruses including SARS-CoV, MERS-CoV, their related enzootic viruses, and endemic common-cold causing coronaviruses (CoV) in various *in vitro* and *in vivo* preclinical models of CoV pathogenesis (Brown et al., 2019; de Wit et al., 2020; Sheahan et al., 2017; Sheahan et al., 2020). More recently, RDV was shown to exert potent antiviral activity against SARS-CoV-2 *in vitro* (Pruijssers et al., 2020) and therapeutic efficacy in a SARS-CoV-2 rhesus macaque model, which recapitulates mild to moderate respiratory symptoms (Williamson et al., 2020). In a double-blind, randomized, placebo-controlled trial (ACTT-1), RDV was shown to shorten recovery time in hospitalized COVID-19 patients by 5 days on average as compared to those receiving placebo (Beigel et al., 2020). In contrast, in an open-label, non-placebo-controlled, and non-blinded clinical trial (WHO Solidarity trial) RDV was not shown to improve outcomes in hospitalized patients (Wang et al., 2020). Importantly, mutations in the viral RNA dependent RNA polymerase (RdRp) known to interfere with the antiviral activity of RDV are not found in the identifying amino acid signatures of SARS-CoV-2 VOCs (Martin et al., 2021). As combinations of RDV with immunomodulators (Baricitinib) have very recently been shown to improve COVID-19 outcomes over single-agent treatment (Kalil et al., 2020), it remains unknown whether RDV combinations with other antiviral drugs with complementary modalities will yield similarly promising results.

Several monoclonal antibodies (mAb) targeting the SARS-CoV-2 spike have been shown to potently neutralize SARS-CoV-2 *in vitro* (Dieterle et al., 2020; Jones et al., 2020; Li et al., 2021; Robbiani et al., 2020; Rogers et al., 2020; Yang et al., 2020; Zost et al., 2020a; Zost et al., 2020b). Monoclonal antibody (mAb) drugs targeting the SARS-CoV-2 spike have demonstrated therapeutic efficacy in multiple pre-clinical models of viral pathogenesis, and a select few have been authorized for emergency use by the FDA to treat COVID-19 (Ly-CoV016/LyCoV555, Eli Lilly; REGN10987/ REGN10933, Regeneron)(2020a; Barnes et al., 2020a; Barnes et al., 2020b; Jones et al., 2020; Schäfer et al., 2021). Most clinical candidate mAbs are RBD-specific and have varying modes of binding and epitope specificities (Barnes et al., 2020a). Lilly’s LY-CoV555 can recognize the RBD in both the up and down conformations (Jones et al., 2020). REGN10987 binds to the RBD outside the ACE2 binding site whereas REGN10933 binds to the top of the RBD and competes with the ACE2 binding site (Hansen et al., 2020). Two recently described highly potent SARS-CoV-2 neutralizing mAbs, C144 and C135, currently being evaluated in human trials at the Rockefeller University Hospital (ClinicalTrials.gov Identifier: NCT04700163) and licensed to Bristol Myers Squibb for development (Robbiani et al., 2020). C144 (IC_50_ = 2.55 ng/mL) and C135 (IC_50_ = 2.98 ng/mL), were isolated from convalescent human patients and target non-overlapping sites on the receptor binding domain (RBD) on the SARS-CoV-2 spike protein similar to the REGN mAb cocktail (Barnes et al., 2020a; Barnes et al., 2020b; Robbiani et al., 2020; Schäfer et al., 2021). As mAb prophylaxis can prevent COVID-19, preliminary results from human clinical trials evaluating the therapeutic efficacy of mAbs in COVID-19 outpatients have thus far been promising (Weinreich et al., 2020; Zhou et al., 2020b).

The emergence of SARS-CoV-2 variants that can partially or completely evade mAbs in advanced clinical development is a growing concern. For example, the SARS-CoV-2 South African B.1.351 variant can completely evade neutralization by mAb LY-CoV555 (Wang et al., 2021a; Wang et al., 2021b). Other mAbs in clinical development, including the AstraZeneca COV2-2196 mAb and the Brii BioSciences mAb Brii-198, have a reduction in neutralization potency by more than 6-fold due to the presence of the E484K mutation (Chen et al., 2021; Wang et al., 2021b). Moreover, the neutralization activity of the Regeneron mAb REGN 10933, is also dampened by the E484K mutation (Wang et al., 2021b). In contrast, the variants do not affect the neutralization potency of C135 (Wang et al., 2021b). Lastly, while the variants do not affect the C144 + C135 antibody combination *in vitro* (Wang et al., 2021c), it is not yet known if this mAb cocktail can protect against the SARS-CoV-2 variants *in vivo*.

We previously developed a mouse-adapted model of SARS-CoV-2 (SARS-CoV-2 MA10) pathogenesis based on the ancestral pandemic strain (Leist et al., 2020). Following SARS-CoV-2 MA10 infection of standard laboratory mice, virus replicates primarily in ciliated epithelial cells and type II pneumocytes with peak titers by 48 hours post infection (hpi) concurrent with body weight loss, loss of pulmonary function, the development of acute lung injury (ALI) and mortality, consistent with severe human COVID-19 pathogenesis (Leist et al., 2020). Here, we define the prophylactic and therapeutic efficacy of RDV and C144 + C135 mAbs used singly and in combination in mice infected with SARS-CoV-2 MA10. We show that the prophylactic and therapeutic administration of RDV or mAb exert a robust antiviral effect and their ability to abrogate disease diminished as a function of initiation time. When combined, RDV/mAb therapy modestly improved outcomes compared to monotherapy suggesting that combination therapy may provide an additional therapeutic benefit over single agents in humans with COVID-19. Importantly, we demonstrate that C144 + C135 mAb combination protects from severe disease against SARS-CoV-2 South African B.1.351 variant challenge in an mouse model of age-related COVID-19 pathogenesis. These data support the continued use of RDV to treat SARS-CoV-2 infections and support the continued clinical development of the C144 and C135 antibody combination to treat patients infected with SARS-CoV-2 variants.

## RESULTS

### Prophylactic and therapeutic RDV protect against COVID-19 disease in mice

First, we sought to determine the time at which RDV therapy would fail to improve outcomes in SARS-CoV-2 infected mice. Due to a serum esterase absent in humans but present in mice that reduces RDV stability (carboxyesterase 1c (*Ces1c*)), we performed all of our RDV efficacy studies in C57BL/6 mice lacking this gene (*Ces1c* ^(-/-)^) (Sheahan et al., 2017). Although we had previously explored the *in vivo* efficacy of RDV against SARS-CoV/SARS-CoV-2 chimeric viruses (Pruijssers et al., 2020), we had not yet evaluated RDV in mice infected with our recently described mouse adapted SARS-CoV-2 (SARS-CoV-2 MA10) (Leist et al., 2020). We initiated twice-daily treatment of mice with a human equivalent dose of RDV (25mg/kg) or vehicle −12 hours prior to infection or 12 (early), 24 (mid-late), or 48 (late) hours post infection (hpi) with 1 × 10^4^ particle forming units (PFU) of SARS-CoV-2 MA10. Body weight loss is a crude marker of emerging coronavirus disease in mice. Body weight loss observed in vehicle treated animals was prevented with prophylactic RDV (Figure 1A). When initiated after SARS-CoV-2 infection, only early therapeutic intervention (+12hr) was able to significantly diminish weight loss (Figure 1A). While RDV therapy initiated at 24hr did not prevent weight loss, lung viral load was significantly diminished in this group similar to those receiving prophylaxis (−12hr) or early therapeutic intervention (+12hr) (Figure 1B). Similarly, lung discoloration, a gross pathologic feature characteristic of severe lung damage, was observed in the vehicle-treated animals but was diminished in all treatment groups except the 48hpi RDV group (Figure 1C). We then used a histologic tool developed by The American Thoracic Society (ATS) to quantitate the pathological features of ALI that we recently utilized to examine the pulmonary pathology of SARS-CoV-2 MA10 infected BALB/c mice (Leist et al., 2020; Matute-Bello et al., 2011). Per animal, three random diseased fields in lung tissue sections were blindly evaluated by a board-certified veterinary pathologist for alveolar septal thickening, protein exudate in the airspace, hyaline membrane formation, and neutrophils in the interstitium or airspaces. Scoring revealed that RDV prophylaxis and therapy initiated at both +12 and +24 hpi reduced ALI as compared to vehicle treated animals (Figure 1D and Figure S1). A complementary histological tool measuring the pathological hallmark of ALI, diffuse alveolar damage (DAD), revealed consistent data (Figure 1E and Figure S1) with those in Figure 1D (Schmidt et al., 2018; Sheahan et al., 2020). Lastly, pulmonary function was measured daily in a subset of mice per group (N = 4) by whole-body plethysmography (WBP). As shown with the WBP metric enhanced pause (PenH), a metric for airway resistance or obstruction that was previously validated in animal models of CoV pathogenesis (Menachery et al., 2015; Sheahan et al., 2017), only prophylactic and early therapeutic administration of RDV (+12hpi) prevented the loss of pulmonary function observed in the other groups. Together, these data show that prophylactic and therapeutic RDV exerts a profound antiviral effect when administered up to 24hpi but the ability of RDV therapy to improve disease outcomes wanes with time of initiation.

**Figure 1.**
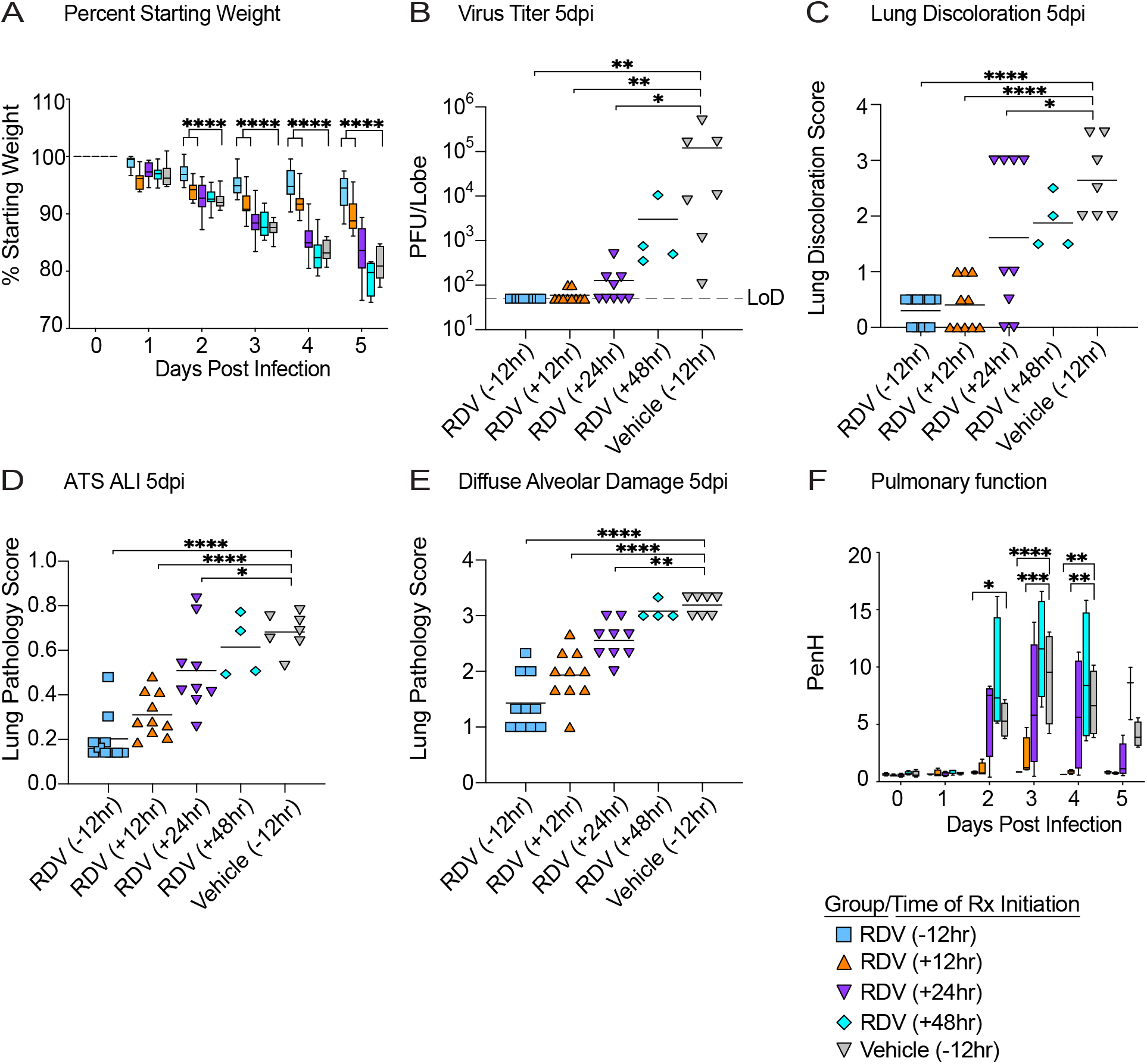
The prophylactic and therapeutic efficacy of RDV against SARS-CoV-2 in mice. (A) % starting weight in prophylactically treated mice with RDV at 12 hours before infection, and therapeutically at 12, 24, and 48 hours post infection. From left to right, light blue bars denote −12 hours prophylactic treatment, orange bars denote +12 hours therapeutic treatment, purple bars denote +24 hours therapeutic treatment, aqua bars denote +48 hours therapeutic treatment, and grey bars denote vehicle treated mice. (B) Lung viral titers in prophylactically and therapeutically treated mice with RDV. Limit of detection (LoD). (C) Lung discoloration score in prophylactically and therapeutically treated mice with RDV. (D-E) Lung pathology in prophylactically and therapeutically treated mice with RDV. (F) Pulmonary function in prophylactically and therapeutically treated mice with RDV. P values are from a 2-way ANOVA after Sidak’s multiple comparisons test.

### Prophylactic and therapeutic single mAb and mAb combinations reduce SARS-CoV-2 pathogenesis

In COVID-19 patients, the time at which mAb therapy loses its protective effect remains unknown. To address this, we sought to determine the prophylactic and therapeutic efficacy of a cocktail of clinical candidate mAbs, C144 and C135, in the SARS-CoV-2 MA10 pathogenesis model noted above. We first established therapeutic efficacy profiles for single mAbs. We treated C57BL/6 mice with mAb C144, mAb C135 or control HIV mAb 12hr before or 12, 24, or 48hr after infection with 1 × 10^4^ PFU of SARS-CoV-2 MA10 (Figure S3, S4, S5 and S6). Both mAbs significantly prevented (prophylactic) or reduced (+12hr, +24hr) SARS-CoV-2 MA10 pathogenesis (body weight loss, lung discoloration, ALI scores, etc.) with C135 exerting more robust protection over C144 with measurable improvements in weight loss and gross pathology even when initiated 48 hpi (Figure S3, S4, and S5). Unlike C135 mAb, C144 mAb did not completely prevent virus replication in the lung when administered at 24hpi suggesting incomplete viral breakthrough (Figure S4) likely driven by mouse adapting Q493K spike mutation which resides in a region critical for C144 binding (Barnes et al., 2020a; Barnes et al., 2020b; Gaebler et al., 2021; Leist et al., 2020). Neither antibody when administered 48hpi could prevent weight loss, lung discoloration or ALI yet viral lung titers were significantly reduced (Figure S6). Together, these data demonstrate that clinical candidate mAb C135 and C144 can both prevent and significantly diminish disease in an ongoing SARS-CoV-2 infection in mice.

Next, we evaluated the prophylactic and therapeutic efficacy of combination C144 + C135 to determine if the single agent therapeutic efficacy could be improved with mAb combinations. Similar to the studies with single agent mAb, we treated C57BL/6 mice with mAb combination C144 + C135 12hr prior to or 12, 24, or 48hr after infection with 1 × 10^4^ PFU of SARS-CoV-2 MA10. Unlike the uniform and consistent body weight loss observed in SARS-CoV-2 MA10 infected mice treated with negative control HIV mAb, prophylactic, early (+12hr) and mid-late (+24hr) therapeutic administration of C144 + C135 mAbs protected against bodyweight loss (Figure 2A). Initiation of therapy 48hpi afforded limited protection from body weight loss (Figure 2A). Remarkably, the levels of infectious virus in the lung were significantly reduced below the limit of detection (50 particle forming units, PFU) in all C144 + C135 mAb groups by day 5 post infection (dpi) unlike control mAb treated animals (mean lung titer = 1×10^4^ PFU/lobe). Mirroring the trend observed in body weight loss, gross lung pathology as measured by observation of lung discoloration was eliminated with prophylactic C144 + C135 mAb, significantly diminished with early (+12hr) and mid-late (+24hr) dosing of C144 + C135 mAb and even moderately reduced with late (+48hr) therapy. We then quantitated the histologic features of ALI using the same tools employed in Figure 1 which demonstrated that prophylactic and therapy initiated up to 24hpi significantly reduced ALI observed in negative control mAb treated animals (Figure 2D). When applying the DAD scoring tool to the same tissue sections, we saw a similar trend yet only prophylactic and early therapeutic (+12hr) C144 + C135 significantly reduced scores (Figure 2E). In agreement with the histological assessment, loss of pulmonary function observed in negative control mAb treated animals could be prevented with prophylactic and early therapeutic (+12hpi) C144 + C135 (Figure 2F).

**Figure 2.**
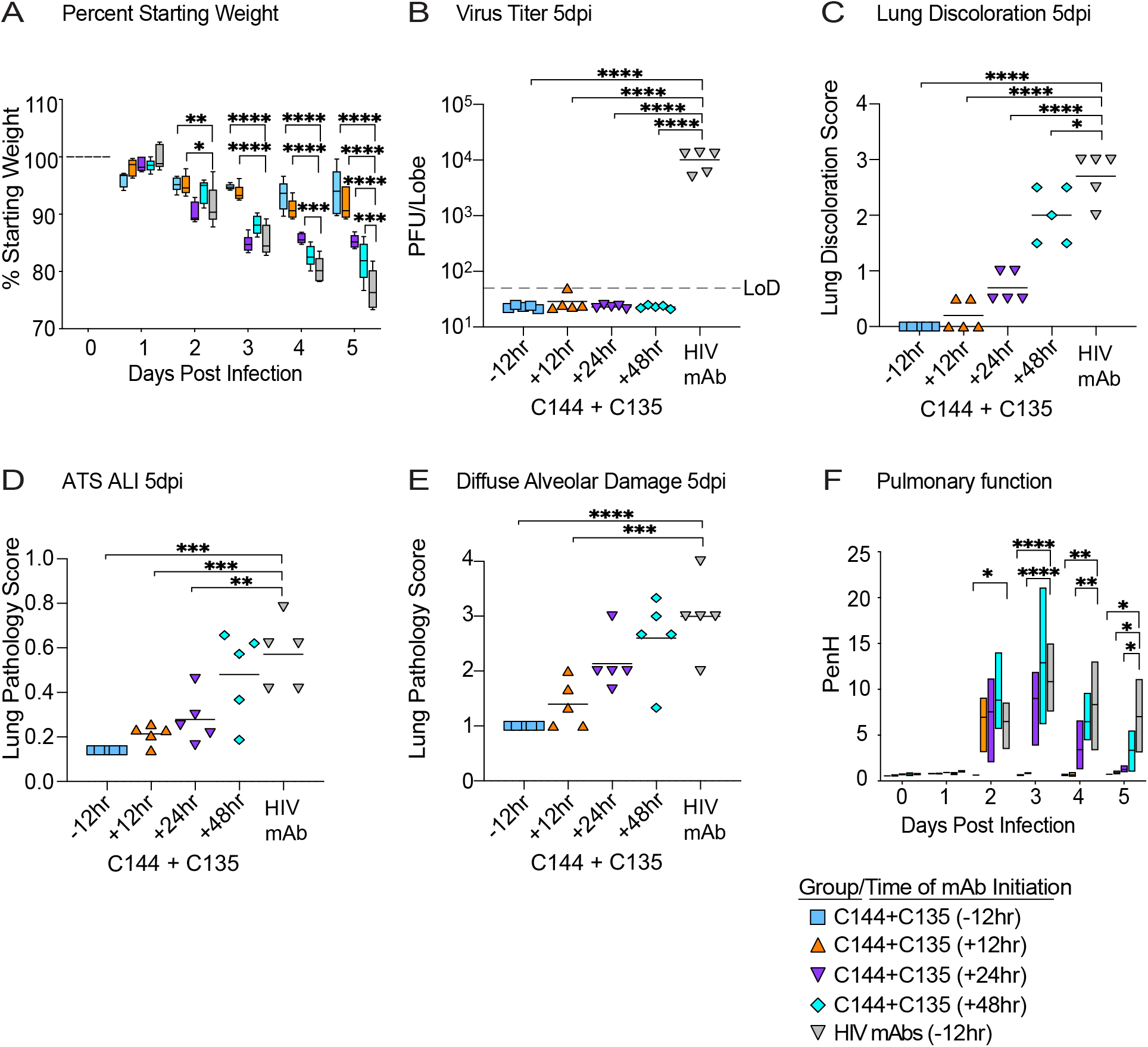
The prophylactic and therapeutic efficacy of mAbs against SARS-CoV-2 in mice. (A) % starting weight in prophylactically treated mice with C144 + C135 at 12 hours before infection, and therapeutically at 12, 24, and 48 hours post infection. From left to right, light blue bars denote −12 hours prophylactic treatment, orange bars denote +12 hours therapeutic treatment, purple bars denote +24 hours therapeutic treatment, aqua bars denote +48 hours therapeutic treatment, and grey bars denote vehicle treated mice. (B) Lung viral titers in prophylactically and therapeutically treated mice with C144 + C135. (C) Lung discoloration score in prophylactically and therapeutically treated mice with C144 + C135. (D-E) Lung pathology in prophylactically and therapeutically treated mice with C144 + C135. (F) Pulmonary function in prophylactically and therapeutically treated mice with C144 + C135. P values are from a 2-way ANOVA after Sidak’s multiple comparisons test.

Interestingly, combination mAb therapy initiated at 24hpi also provided a benefit in pulmonary function (Figure 2F). Thus, mAb therapy can exert a profound antiviral effect even when administered at later times post infection.

### Combination RDV/mAb cocktail demonstrates a small improvement vs mAb therapy alone at 36hpi

We sought to determine if combination RDV/C144+C135 mAb would further curtail viral pathogenesis over that provided by single agents. We designed a study where we initiated single agent or combination therapy 24hr after SARS-CoV-2 MA10 infection, treated mice up to 7dpi and followed mice until 12dpi to determine if therapy accelerated recovery. Among groups receiving single agents or combination therapies, significant differences in body weight were not consistently noted (Figure S7A) but all therapeutic treatment groups provided complete protection from mortality observed with vehicle treatment (Figure S7B). Upon completion of the study on 12dpi, differences in gross pathology were not noted among treatment groups (Figure S7C). We performed pulmonary function by WBP on select groups (i.e. vehicle/control mAb and RDV/mAb combination) for the first 5 days of infection and observed a rapid improvement in pulmonary function with combination therapy which returned to baseline by 3dpi (Figure S7D).

To determine if a further delay treatment initiation time closer to peak of virus replication in the lung would reveal an improved benefit of combination therapy, we performed a therapeutic efficacy study initiating treatment at 36hpi. Rather than focus on the potential effects on recovery, the goal of this study was to determine if combination therapy had a differential effect on lung pathology and virus replication during the acute phase of disease. We initiated treatment 36hr after infection with 1 × 10^4^ PFU SARS-CoV-2 MA10 in C57BL/6 (*Ces1c* ^(-/-)^) mice with the vehicle, single agent, and combination groups as described in the previous combination experiment. We observed a small but measurable improvement in body weight loss with RDV/mAb treatment (Figure 3A). Similarly, by 3dpi, only the RDV/control mAb and RDV/mAb-treated groups had lower lung viral titers compared to the vehicle/control mAb-treated group (Figure 3B). By 5dpi, vehicle treated animals had mean lung titers nearing 1 x 10^5^ PFU, yet all treatment groups had significantly reduced lung titers at or near the limit of detection (Figure 3C). When examining gross lung pathology 5dpi, all therapies provided significant protection from lung discoloration observed with vehicle treatment, but RDV/mAb combination therapy group had the overall lowest score and was significantly improved over single agent vehicle/mAb (Figure 3D). We then quantitated the histological manifestations of ALI using the two complementary scoring tools employed above. With both ATS and DAD scoring systems, ALI was readily apparent in vehicle treated animals (Figure 3E and 3F). Although mirroring the trend observed in the gross pathological observations where combination therapy afforded protection over single agent therapy, significant differences were not observed among groups receiving antiviral therapies and all reduced ALI on 5dpi (Figure 3E and 3F). Lastly, we examined the effect of combination therapy on pulmonary function (Figure 3G). Combination RDV/mAb initiated at 36hpi reduced the loss of pulmonary function observed with vehicle treatment on 3-5dpi (Figure 3G). Altogether, our findings suggest that combination therapy with RDV and potent neutralizing mAbs provides a small but measurable benefit over single agents in some but not all metrics of SARS-CoV-2 pathogenesis in this model.

**Figure 3.**
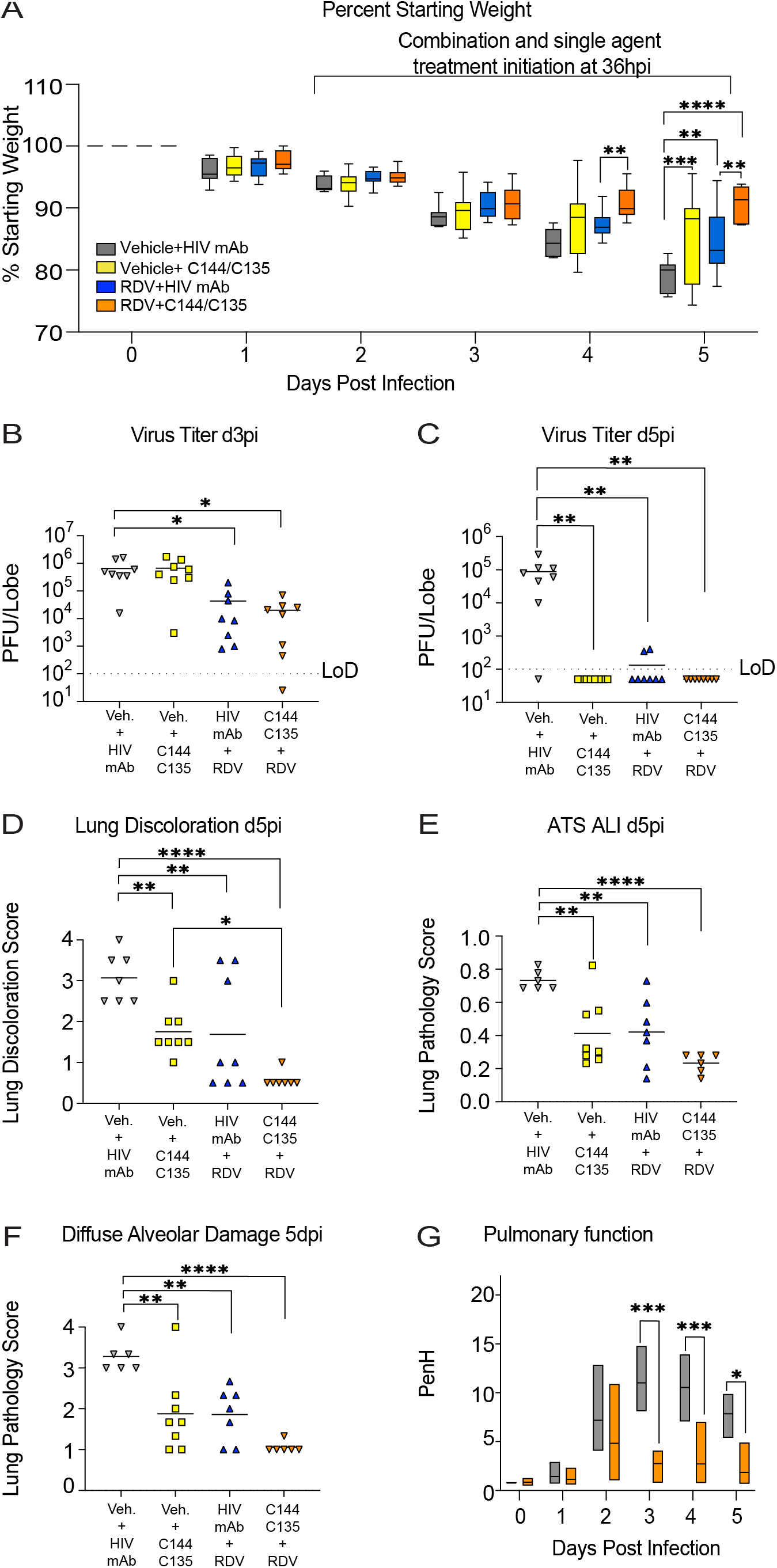
The therapeutic efficacy of RDV and mAbs as single agents and in combination at 36 hours post infection in SARS-CoV-2-infected mice. (A) % starting weight in therapeutically treated mice with vehicle + HIV mAb, vehicle + C144 + C135, RDV + HIV mAb, and RDV + C144 + C135 at 36 hours post infection. From left to right, grey bars denote vehicle/control mAb treated mice, yellow bars denote vehicle/mAb therapeutic treatment, blue bars denote RDV/control mAb therapeutic treatment, and orange bars denote RDV/mAb therapeutic treatment. (B) Day 3 post infection lung viral titers in therapeutically treated mice with single agents and combination therapy. “Veh.” signifies vehicle treatment. (C) Day 5 post infection lung viral titers in therapeutically treated mice with single agents and combination therapy. (D) Lung discoloration scores in therapeutically treated mice with single agents and combination therapy. (E) Pulmonary function in therapeutically treated mice with vehicle + HIV mAb and RDV + C144 + C135. P values are from a 2-way ANOVA after Sidak’s multiple comparisons test.

### C144+C135 mAb prophylaxis and therapy improve outcomes in South African B.1.351 variant of concern infected mice

The emergence of neutralization-resistant SARS-CoV-2 variants is a growing threat. B.1.351, which initially emerged in South Africa, is a VOC that can infect mice without adaptation (Montagutelli et al., 2021). B.1.351 has characteristic RBD mutations at residues K417, E484, and N517 which result in resistance to many of the class 1 and 2 antibodies that dominate the initial RBD-directed neutralizing response (Barnes et al., 2020a; Chen et al., 2021; Planas et al., 2021; Wang et al., 2021c). For example, B.1.351 is completely resistant to Eli Lilly’s Ly-CoV555 mAb (Wang et al., 2021a), underlining the importance of monitoring the *in vivo* efficacy of monoclonal antibody therapies that are in advanced clinical testing against SARS-CoV-2 VOCs. To examine the *in vivo* efficacy of the C144 + C135 mAb combination against recombinant mouse adapted SARS-CoV-2 bearing the B.1.351 spike, we treated aged BALB/c mice with mAb 12hr before or after infection with 1 × 10^4^ PFU. Weight loss observed with control antibody treatment was prevented with C144 + C135 prophylaxis and lung viral loads were reduced below the limit of detection on both 3 and 5dpi (Figure 4A-C). Similarly, mAb combination therapy accelerated recovery and diminished virus replication below the limit of detection by 5dpi (Figure 4A and 4C). To complement the infectious virus data, we then quantitated viral subgenomic RNAs in mouse lung tissues in each group. Unlike the quantitation of SARS-CoV-2 genomic RNA, which has the potential to measure RNA from infectious particles, defective particles, mAb bound particles and various replicative forms of viral RNA, these subgenomic RNA qRT-PCR assays are specific for envelope (E) and nucleocapsid (N) viral transcripts which are only made in actively replicating cells. Prophylactic and therapeutic administration of C144 + C135 significantly reduced lung viral E sgRNA (Figure 4D and 4E) and N sgRNA (Figure 4F and 4G) compared to the control mAb treated animals indicating that mAb therapy successfully reduced levels of replication of SARS-CoV-2 bearing the B.1.351 spike *in vivo*. Finally, gross pathology caused by mouse adapted SARS-CoV-2 bearing the B.1.351 spike was significantly reduced in aged mice with both prophylactic and therapeutic administration of the C144 + C135 combination (Figure 4H and 4I). Collectively, these data demonstrate that both prophylaxis and therapy with combination C144 + C135 mAb can potently reduce virus replication and improve disease outcomes *in vivo* following infection with variant B.1.351.

**Figure 4.**
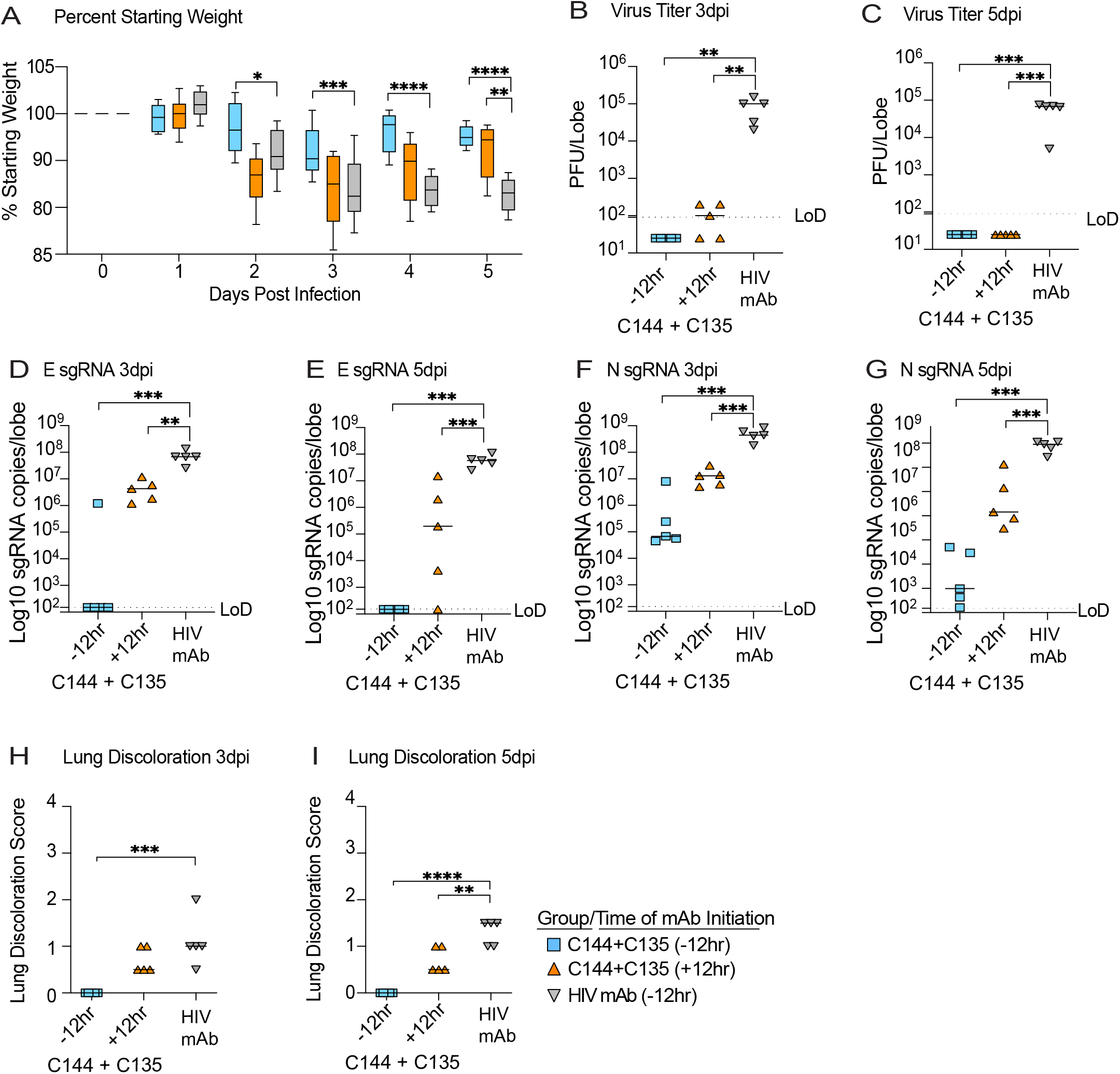
The prophylactic and therapeutic efficacy of C144 + C135 against SARS-CoV-2 B.1.351 in aged mice. (A) % starting weight in prophylactically treated mice with C144 + C135 at 12 hours before infection, and therapeutically at 12 post infection. From left to right, light blue bars denote −12 hours prophylactic treatment, orange bars denote +12 hours therapeutic treatment, and grey bars denote prophylactically treated mice with HIV mAb. (B-C) Lung viral titers at day 3 and 5 post infection in prophylactically and therapeutically treated mice with C144 + C135 and HIV mAb negative controls. (D-E) Sugenomic Envelope (E) RNA copies/lobe in prophylactically and therapeutically treated mice with C144 + C135 and HIV mAb. (F-G) Sugenomic Envelope (N) RNA copies/lobe in prophylactically and therapeutically treated mice with C144 + C135 and HIV mAb. (H-I) Lung discoloration at day 3 and 5 post infection in prophylactically and therapeutically treated mice with C144 + C135 and HIV mAb. P values are from a 1-way ANOVA following Dunnett’s multiple comparisons.

## DISCUSSION

Therapies effective against the current and future SARS-CoV-2 VOCs are desperately needed to treat those yet to be vaccinated or those experiencing breakthrough infection. RDV is a broad-spectrum antiviral drug and has potent antiviral activity against multiple emerging, endemic and enzootic CoVs including: SARS-CoV, SARS-CoV-2, MERS-CoV, bat-CoV WIV-1, bat-CoV RsSHC014, bat-CoV HKU5, bat-CoV HKU-3-1, HCoV-229, HCoV-NL63, HCoV-OC43, porcine deltacoronavirus (PDCoV) (Agostini et al., 2018; Brown et al., 2019; de Wit et al., 2020; Sheahan et al., 2017). In addition to the *in vitro* activity of RDV against SARS-CoV-2 (Pruijssers et al., 2020), RDV can exert an antiviral effect and diminish SARS-CoV-2 disease in rhesus macaques which develop mild respiratory disease (Williamson et al., 2020). Similarly, the prophylactic efficacy of mAb C144 and C135 have previously been evaluated in replication models of mouse adapted SARS-CoV-2 based on the ancestral pandemic strain (Schäfer et al., 2021), but their prevention and therapy has not yet been evaluated in the context of the emerging variants that can evade vaccine-elicited antibodies and existing mAb therapies.

Human clinical data for direct antivirals like mAb and small molecule antivirals like RDV provides clear evidence that their success at improving outcomes is directly related to the time after the onset of symptoms that therapy is initiated. Outpatient studies evaluating mAb drugs in humans with mild to moderate COVID-19 demonstrated notable reductions in virus shedding and symptoms, which enabled the FDA emergency use authorization (EUA) of both Eli Lilly’s and Regeneron’s antibody cocktails (Gottlieb et al., 2021; Weinreich et al., 2020). However, hospitalized patients with advanced COVID-19 disease treated with these mAb drugs did not have measurably improved outcomes compared to standard of care (2020b). While RDV has been shown to accelerate recovery of COVID-19 hospitalized patients (Beigel et al., 2020), insight in to whether RDV will further improve outcomes in patients earlier in the course of COVID-19 remains unknown. Thus, the optimal window after the onset of symptoms within which to treat with antivirals such as RDV or potent mAbs fail remains unknown.

In this manuscript, we aimed to define the time after SARS-CoV-2 infection in mice where RDV or mAb therapy fail to exert an antiviral effect and/or fail to improve disease outcomes. Like mouse-adapted models of SARS-CoV and MERS-CoV, the replication kinetics of mouse-adapted SARS-CoV-2 MA10 in mice is compressed with peak replication in the lung 48hpi (Leist et al., 2020). In contrast, the replication kinetics of SARS-CoV-2 in the airways of humans is more variable with reports estimating peak replication within the first week after the onset of symptoms (Liu et al., 2020; Zheng et al., 2020). Moreover, human patients can shed viral RNA in the mucosa of the upper respiratory tract as long as 24 days post infection (Zhou et al., 2020a), underlining that sustained viral shedding and symptoms can last considerably longer in humans than mice. Thus, the window within which to intervene with antiviral therapy prior to the peak of virus replication in humans is dramatically different than in mice (~2 days). While our mouse model faithfully recapitulates many aspects of human COVID-19 disease (e.g. high titer replication in the upper and lower airway, loss of pulmonary function, acute lung injury, age related exacerbation of disease, etc.), it is not possible to very finely correlate the compressed kinetics of disease in mouse and those in humans but there are a few notable takeaways from the modeling presented herein. Given early therapeutic treatment at +12 and +24hpi in our model provided the most benefit, it is likely the benefit of antibody and small molecule antivirals like RDV will be maximized if given prior to peak viral replication and/or early in the disease course before patients are hospitalized. In addition, we show a small improvement with combination mAb/RDV over single agent therapy which suggests that combinations of antiviral drugs of disparate modalities may offer an additional benefit in COVID-19 patients over single agents, something that should be rigorously evaluated in humans. Although our studies clearly support the use/evaluation of RDV and mAb as treatments for COVID-19, both are administered intravenously limiting their broad distribution to COVID-19 outpatients. Potential strategies to allow the wider dissemination of these treatments may include chemical alteration of RDV to facilitate oral bioavailability and/or less complicated subcutaneous or intramuscular injections of mAbs. The effect of mAb injection route (i.e. subcutaneous vs. intravenous) on pharmacokinetics and safety is currently being evaluated for C144 and C135 in Phase I clinical studies (ClinicalTrials.gov Identifier: NCT04700163).

Given the growing emergence of SARS-CoV-2 variants, we examined the prophylactic and therapeutic efficacy of the C144 + C135 combination against the South African B.1.351 variant spike in a robust age-related mouse model of SARS-CoV-2 pathogenesis. Importantly, the C144 + C135 cocktail demonstrated prophylactic and therapeutic efficacy against the B.1.351 VOC, which is encouraging given that this variant has demonstrated full escape from other mAbs approved for emergency use in humans, such as the LY-CoV555. In addition, the neutralizing potency of the AstraZeneca and Brii Biosciences mAbs in clinical trials are clearly dampened by mutations present in the variants such as the B.1.351 (Wang et al., 2021b). The target of the antiviral activity of RDV is the viral RdRp. Importantly, hallmark mutations of current SARS-CoV-2 VOCs are not found in regions of the RdRp known to affect the antiviral potency of RDV, thus antiviral resistance to RDV is not currently anticipated with current VOCs (Martin et al., 2021). In context of emerging variants in the future, it will be critical to continue to evaluate the prevention and therapy of currently approved small molecule and mAb antivirals and those in clinical development against newly emerging variants of interest. Our results reveal that prophylaxis and therapy with the C144 + C135 mAb combination is robustly antiviral against the B.1.351 VOC spike *in vivo* and can diminish the development of disease during an ongoing SARS-CoV-2 infection in mice. These data support the further evaluation of this mAb cocktail as therapy in human patients infected with the B1.351 variant.

## STAR METHODS

### Lead Contact

Further information and requests for resources and reagents should be directed to and will be fulfilled by the Lead Contact, Timothy P. Sheahan

### Materials Availability

Not applicable.

### Data and Code availability

Not applicable.

## EXPERIMENTAL MODEL AND SUBJECT DETAILS

### Animals and virus infections

Twenty-week-old male and female *Ces1c* ^(-/-)^ on a B6 background (C57BL/6J: Jackson Laboratory # 014096) were purchased from Jackson Laboratory. Eleven-month-old female BALB/c mice were purchased from Envigo (#047). A mouse-adapted SARS-CoV-2 virus (MA10) was used in all experiments and this virus was previously described (Leist et al., 2020). Briefly, mutations predictive of increased affinity to mouse ACE2 were introduced into a SARS-CoV-2 virus plasmid system and the virus was recovered by reverse genetics (Dinnon et al., 2020). This modified virus was then serially passaged in aged BALBc mice (Envigo #047) for ten passages which we refer to as the mouse-adapted passage 10 (MA10) SARS-CoV-2 (Leist et al., 2020). A mouse-adapted (MA10) backbone expressing the SARS-CoV-2 B.1.351 spike was generated for this study. All mice were anesthetized and infected with SARS-CoV-2 MA10 or B.1.351 spike/MA10 intranasally with 1 × 10^4^ PFU/ml. Mice were weighed daily and were monitored for signs of SARS-CoV-2 clinical disease in all experiments.

### Animal care

The study was carried out in accordance with the recommendations for care and use of animals by the Office of Laboratory Animal Welfare (OLAW), National Institutes of Health and the Institutional Animal Care and Use Committee (IACUC) protocol number: 20-059 at University of North Carolina (UNC permit no. A-3410-01). Virus inoculations were performed under anesthesia and all efforts were made to minimize animal suffering. Animals were housed in groups and fed standard chow diets.

## METHOD DETAILS

### Study design and treatment groups

For the RDV experiment, a total of N=40, ~20-week-old male and female mice were divided into four groups each with N=10 mice with equal numbers of females and males in each group. RDV was administered subcutaneously twice per day (BID) at 25 mg/kg. Groups of N=10 mice (N=5 males and N=5 females) were used in either the prophylaxis −12 hours before infection group, the early therapeutic 12 hours post infection group, the mid-late therapeutic 24 hours post infection group, and the late therapeutic 48 hours post infection group.

For the initial monoclonal antibody experiment, mice were infected as described above and weighed daily and were monitored for signs of SARS-CoV-2 clinical disease. A total amount of 200 μg of C144 + C135, 200 μg of C144, 200 μg of C135, and 200 μg of HIV mAbs 3BC117 + 10-1074 was administered intraperitonially once by injection for each intervention group. Groups of N=20 female mice (N=5 mice treated with C144 + C135, N=5 mice treated with C144, N=5 mice treated with C135, and N=5 mice treated with 3BNC117 + 10-1074) were administered antibody 12 hours before infection, N=20 female mice (N=5 mice treated with C144 + C135, N=5 mice treated with C144, N=5 mice treated with C135, and N=5 mice treated with 3BNC117 + 10-1074) were administered antibody 12hpi (early therapeutic group), N=20 female mice (N=5 mice treated with C144 + C135, N=5 mice treated with C144, N=5 mice treated with C135, and N=5 mice treated with 3BNC117 + 10-1074) were administered antibody 24hpi (mid-late therapeutic group), and N=20 female mice (N=5 mice treated with C144 + C135, N=5 mice treated with C144, N=5 mice treated with C135, and N=5 mice treated with 3BNC117 + 10-1074) were administered antibody 48hpi (late therapeutic group).

For the 24hpi drug and mAb combination intervention experiment, a total of N=40, ~20-week-old male and female mice were divided into four groups each with N=10 mice with equal numbers of females and males in each group. At 24hpi, RDV treatment was initiated by subcutaneous injection twice per day (BID) at 25 mg/kg, and a total amount of 200 μg of C144 + C135 was administered intraperitonially once by injection. N=10 mice (N=5 males and N=5 females) were used in the vehicle + HIV mAb group. N=10 mice (N=5 males and N=5 females) were used in the vehicle + C144 + C135 mAb group. N=10 mice (N=5 males and N=5 females) were used in the RDV + HIV mAb group. N=10 mice (N=5 males and N=5 females) were used in the RDV + C144 + C135 mAb group.

For the 36hpi drug + mAb combination intervention experiment, a total of N=64, ~20-week-old male and female mice were divided into four groups each with N=16 mice with an equal number of females and males in each group. At 36hpi, RDV treatment was initiated by subcutaneous injection twice per day (BID) at 25 mg/kg, and a total of 200 μg of each monoclonal antibody treatment was administered intraperitonially once by injection. N=32 mice were harvested at d3pi to evaluate early lung viral replication titers, and remaining mice were harvested at d5pi. N=16 mice (N=8 males and N=8 females) were used in the vehicle + HIV mAb group. N=16 mice (N=8 males and N=8 females) were used in the vehicle + C144 + C135 mAb group. N=16 mice (N=8 males and N=8 females) were used in the RDV + HIV mAb group. N=16 mice (N=8 males and N=8 females) were used in the RDV + C144 + C135 mAb group.

### Lung pathology scoring

Acute lung injury was quantified via two separate lung pathology scoring scales: Matute-Bello and Diffuse Alveolar Damage (DAD) scoring systems. Analyses and scoring were performed by a Board Certified Veterinary Pathologist who was blinded to the treatment groups as described previously (Sheahan et al., 2020). Lung pathology slides were read and scored at 600X total magnification.

The lung injury scoring system used is from the American Thoracic Society (Matute-Bello) in order to help quantitate histological features of ALI observed in mouse models to relate this injury to human settings. In a blinded manner, three random fields of lung tissue were chosen and scored for the following: (A) neutrophils in the alveolar space (none = 0, 1–5 cells = 1, > 5 cells = 2), (B) neutrophils in the interstitial septae (none = 0, 1–5 cells = 1, > 5 cells = 2), (C) hyaline membranes (none = 0, one membrane = 1, > 1 membrane = 2), (D) Proteinaceous debris in air spaces (none = 0, one instance = 1, > 1 instance = 2), (E) alveolar septal thickening (< 2x mock thickness = 0, 2–4x mock thickness = 1, > 4x mock thickness = 2). To obtain a lung injury score per field, A–E scores were put into the following formula score = [(20x A) + (14 x B) + (7 x C) + (7 x D) + (2 x E)]/100. This formula contains multipliers that assign varying levels of importance for each phenotype of the disease state. The scores for the three fields per mouse were averaged to obtain a final score ranging from 0 to and including 1.

The second histology scoring scale to quantify acute lung injury was adopted from a lung pathology scoring system from lung RSV infection in mice (Schmidt et al., 2018). This lung histology scoring scale measures diffuse alveolar damage (DAD). Similar to the implementation of the ATS histology scoring scale, three random fields of lung tissue were scored for the following in a blinded manner: 1= absence of cellular sloughing and necrosis, 2=Uncommon solitary cell sloughing and necrosis (1–2 foci/field), 3=multifocal (3+foci) cellular sloughing and necrosis with uncommon septal wall hyalinization, or 4=multifocal (>75% of field) cellular sloughing and necrosis with common and/or prominent hyaline membranes. The scores for the three fields per mouse were averaged to get a final DAD score per mouse. The microscope images were generated using an Olympus Bx43 light microscope and CellSense Entry v3.1 software.

### Remdesivir (RDV)

RDV was synthesized at Gilead Inc., and its chemical composition and purity were analyzed by nuclear magnetic resonance, high resolution mass spectrometry, and high-performance liquid chromatography. RDV was solubilized in 12% sulfobutylether-β-cyclodextrin in water (with HCl/NaOH) at pH 5 for *in vivo* studies in mice. RDV was made available to UNC Chapel Hill under an existing material transfer agreement with Gilead Sciences Inc.

### RNA extraction and subgenomic RNA assay

Lung lobes were harvested and homogenized in 1ml of TRIzol reagent. RNA was extracted with phenol/chloroform/isoamyl alcohol solution (25:24:1), precipitated with isopropyl alcohol, washed with 75% ethanol, and resuspended in RNAase-free water. SARS-CoV-2 E gene and N gene subgenomic mRNA (sgRNA) was measured by a one-step RT-qPCR adapted from previously described methods (Li et al., 2021). RNA extracted from animal samples or RNA standards were then measured using TaqMan Fast Virus 1-Step Master Mix (ThermoFisher, catalog # 4444432) and custom primers/probes targeting the E gene sgRNA (forward primer: 5’ CGATCTCTTGTAGATCTGTTCTCE 3’; reverse primer: 5’ ATATTGCAGCAGT ACGCACACA 3’; probe: 5’ FAM ACACTAGCCATCCTTACTGCGCTTCG-BHQ1 3’) or the N gene sgRNA (forward primer: 5’ CGATCTCTTGTAGATCTGTTCTC 3’; reverse primer: 5’ GGTGAA CCAAGACGCAGTAT 3’; probe: 5’ FAM-TAACCAGAATGGAGAACGCAGTG GG-BHQ1 3’). RT-QPCRreactions were carried out on a CFX Opus 384 machine (Bio-Rad) using a program below: reverse transcription at 50°C for 5 minutes, initial denaturation at 95°C for 20 seconds, then 40 cycles of denaturation-annealing-extension at 95°C for 15 seconds and 60°C for 30 seconds. Standard curves were used to calculate E or N sgRNA in copies per ml; the limit of detections (LOD) for both E and N sgRNA assays were 150 copies per lung lobe.

### Biocontainment and biosafety

Studies were approved by the UNC Institutional Biosafety Committee approved by animal and experimental protocols in the Baric laboratory. All work described here was performed with approved standard operating procedures for SARS-CoV-2 in a biosafety level 3 (BSL-3) facility conforming to requirements recommended in the Microbiological and Biomedical Laboratories, by the U.S. Department of Health and Human Service, the U.S. Public Health Service, and the U.S. Center for Disease Control and Prevention (CDC), and the National Institutes of Health (NIH).

### Statistics

All statistical analyses were performed using GraphPad Prism 9. Statistical tests used in each figure are denoted in the corresponding figure legend. A Sidak’s multiple comparisons test was used following 2-way ANOVAs and this is also denoted in the figure legends.

## Acknowledgements

David R. Martinez is funded by a Burroughs Wellcome Fund Postdoctoral Enrichment Program Award, a Hanna H. Gray Fellowship from the Howard Hughes Medical Institute, and was supported by an NIH NIAID T32 AI007151 and an NIH F32 AI152296. This work was supported by NIAID R01 AI132178 awarded to Timothy P. Sheahan and Ralph S. Baric, and an NIH animal models contract (HHSN272201700036I) to R.S.B. This project was also supported by the North Carolina Policy Collaboratory at the University of North Carolina at Chapel Hill with funding from the North Carolina Coronavirus Relief Fund established and appropriated by the North Carolina General Assembly. Michel C. Nussenzweig is an investigator of the Howard Hughes Medical Institute. Animal histopathology services were performed by the Animal Histopathology & Laboratory Medicine Core at the University of North Carolina, which is supported in part by an NCI Center Core Support Grant (5P30CA016086-41) to the UNC Lineberger Comprehensive Cancer Center.

## Author contributions

Conceived the study: D.R.M, A.S., R.S.B, M.C.N., and T.P.S. Designed experiments: D.R.M, A.S., J.Y.F., E.B., D.P.P., T.C., R.S.B, M.C.N., and T.P.S. Performed laboratory experiments: D.R.M, A.S., S.R. L., D.L., K.G., T.P.S.; Provided critical reagents: J.Y.F., E.B., D.P.P., T.C.; Wrote the first draft of the paper: D.R.M and T.P.S. Edited the manuscript: D.R.M, A.S., S.R. L., K.G., J.Y.F., E.B., D.P.P., T.C., S.A.M., B.F.H., R.S.B, M.C.N., and T.P.S. All authors reviewed and approved the manuscript.

## Supplemental figure legends

**Figure S1.**
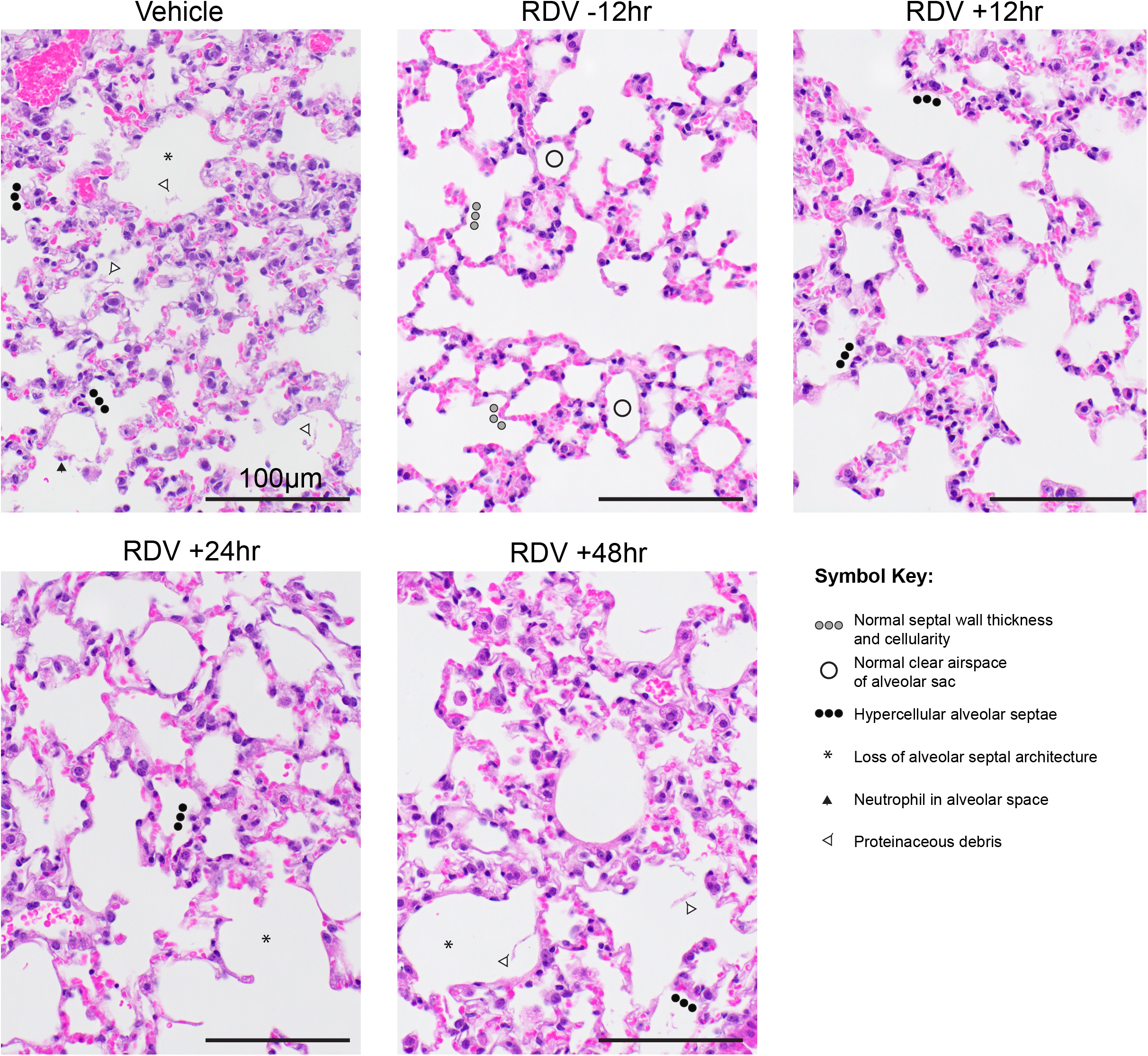
Lung pathology of SARS-CoV-2-infected mice treated with RDV and vehicle prophylactically and therapeutically. Pathologic features of acute lung injury were scored using two separate tools: the American Thoracic Society Lung Injury Scoring (ATS ALI) system. Using this ATS ALI system, we created an aggregate score for the following features: neutrophils in the alveolar and interstitial space, hyaline membranes, proteinaceous debris filling the air spaces, and alveolar septal thickening. Three randomly chosen high power (×60) fields of diseased lung were assessed per mouse. Representative images are shown from vehicle and RDV-treated mice. Symbols identifying example features of disease are indicated in the figure. All images were taken at the same magnification. The black bar indicates 100 μm scale.

**Figure S2.**
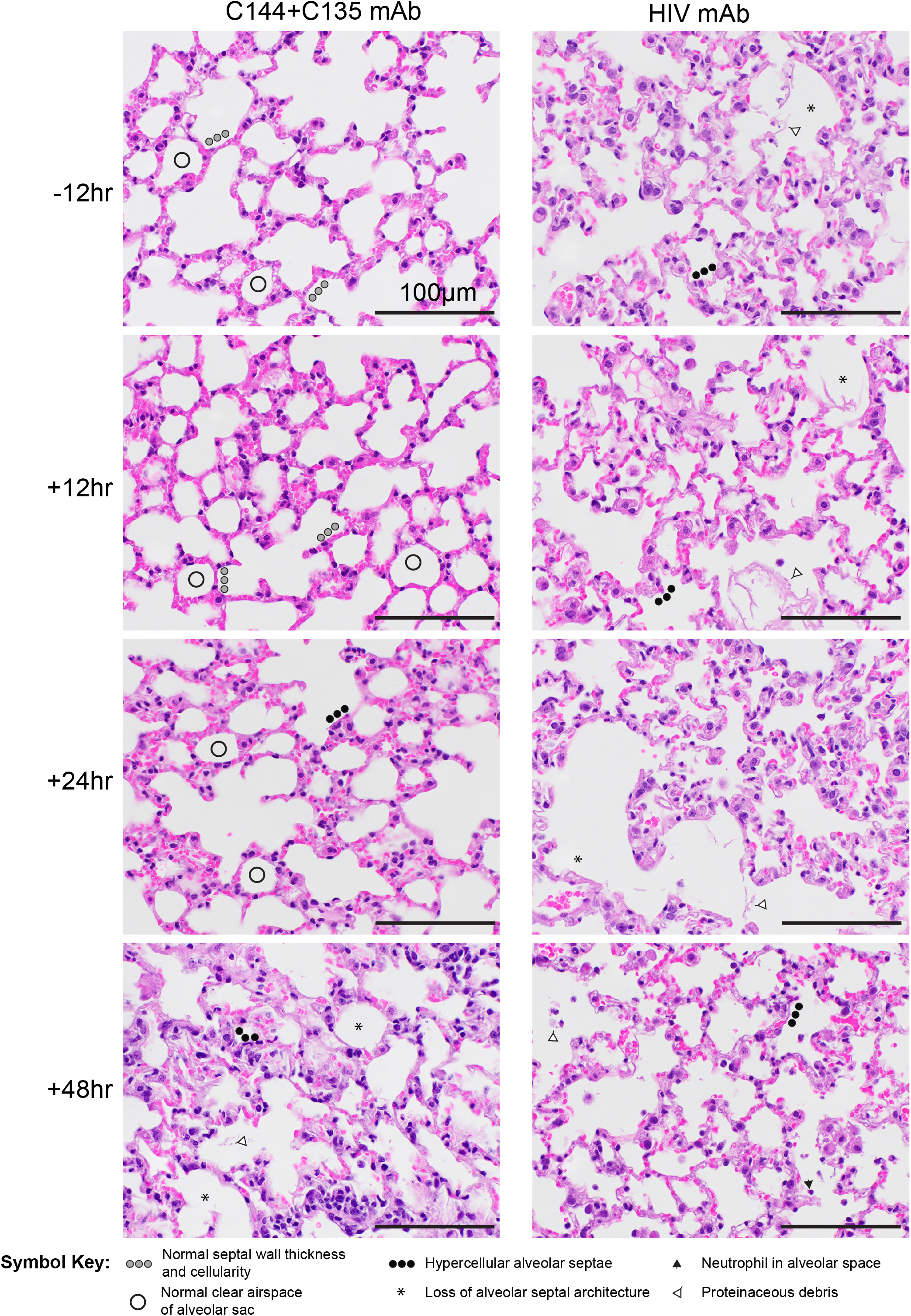
Lung pathology of SARS-CoV-2-infected mice treated with C144 + C135 and an HIV mAb prophylactically and therapeutically. Pathologic features of acute lung injury were scored using two separate tools: the American Thoracic Society Lung Injury Scoring (ATS ALI) system. Using this ATS ALI system, we created an aggregate score for the following features: neutrophils in the alveolar and interstitial space, hyaline membranes, proteinaceous debris filling the air spaces, and alveolar septal thickening. Three randomly chosen high power (×60) fields of diseased lung were assessed per mouse. Representative images are shown from HIV mAb and C144 + C135-treated mice. Symbols identifying example features of disease are indicated in the figure. All images were taken at the same magnification. The black bar indicates 100 μm scale.

**Figure S3.**
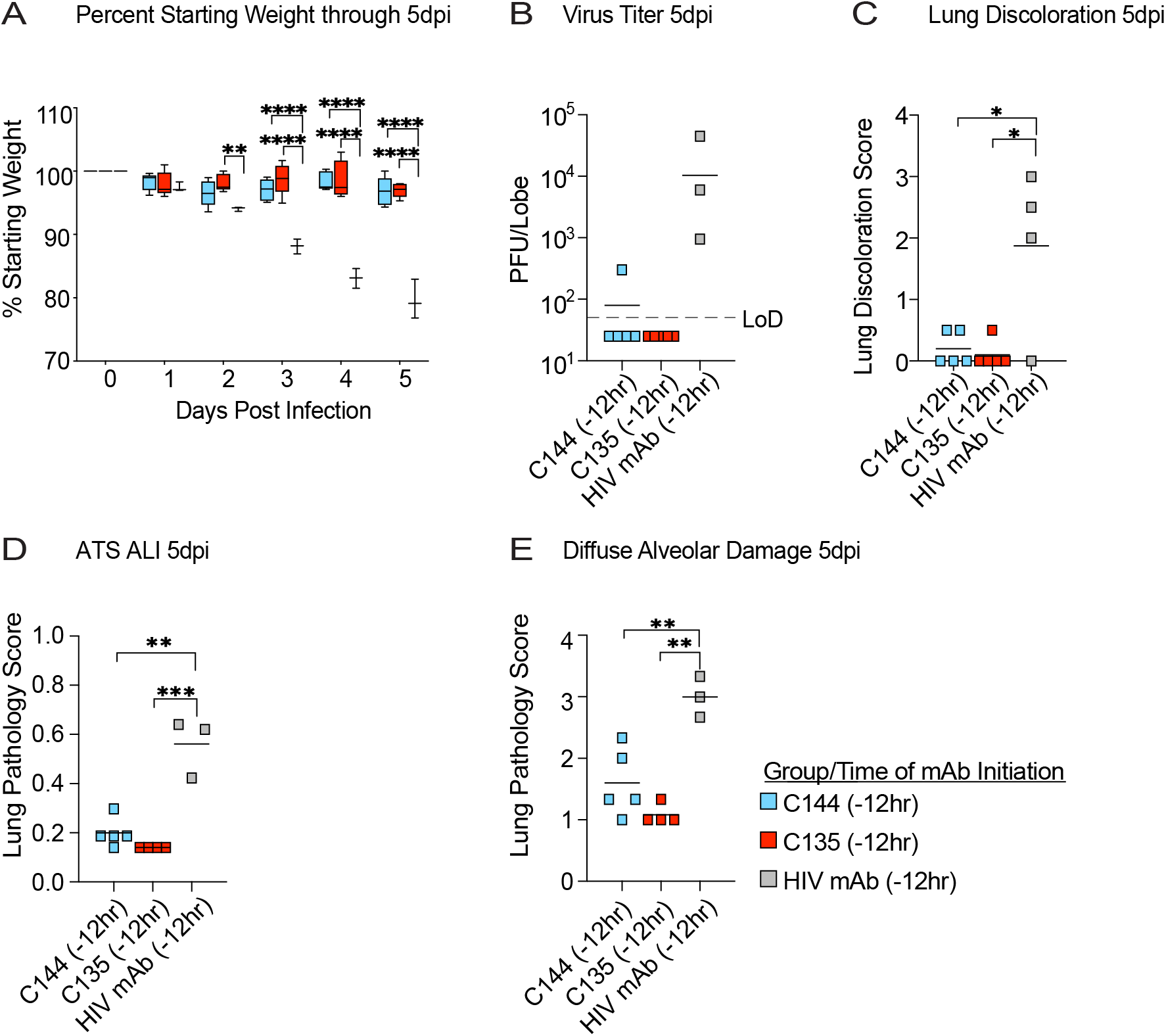
The prophylactic efficacy of mAb monotherapy against SARS-CoV-2 in mice treated at 12 hours before infection. (A) % starting weight in therapeutically treated mice with C144, C135, or an HIV mAb at 12 hours before infection. (B) Lung viral titers in therapeutically treated mice at 12 hours before infection. (C) Lung discoloration score in therapeutically treated mice at 12 hours before infection. (D-E) Lung pathology in therapeutically treated mice at 12 hours before infection. P values are from a 2-way ANOVA after Sidak’s multiple comparisons test.

**Figure S4.**
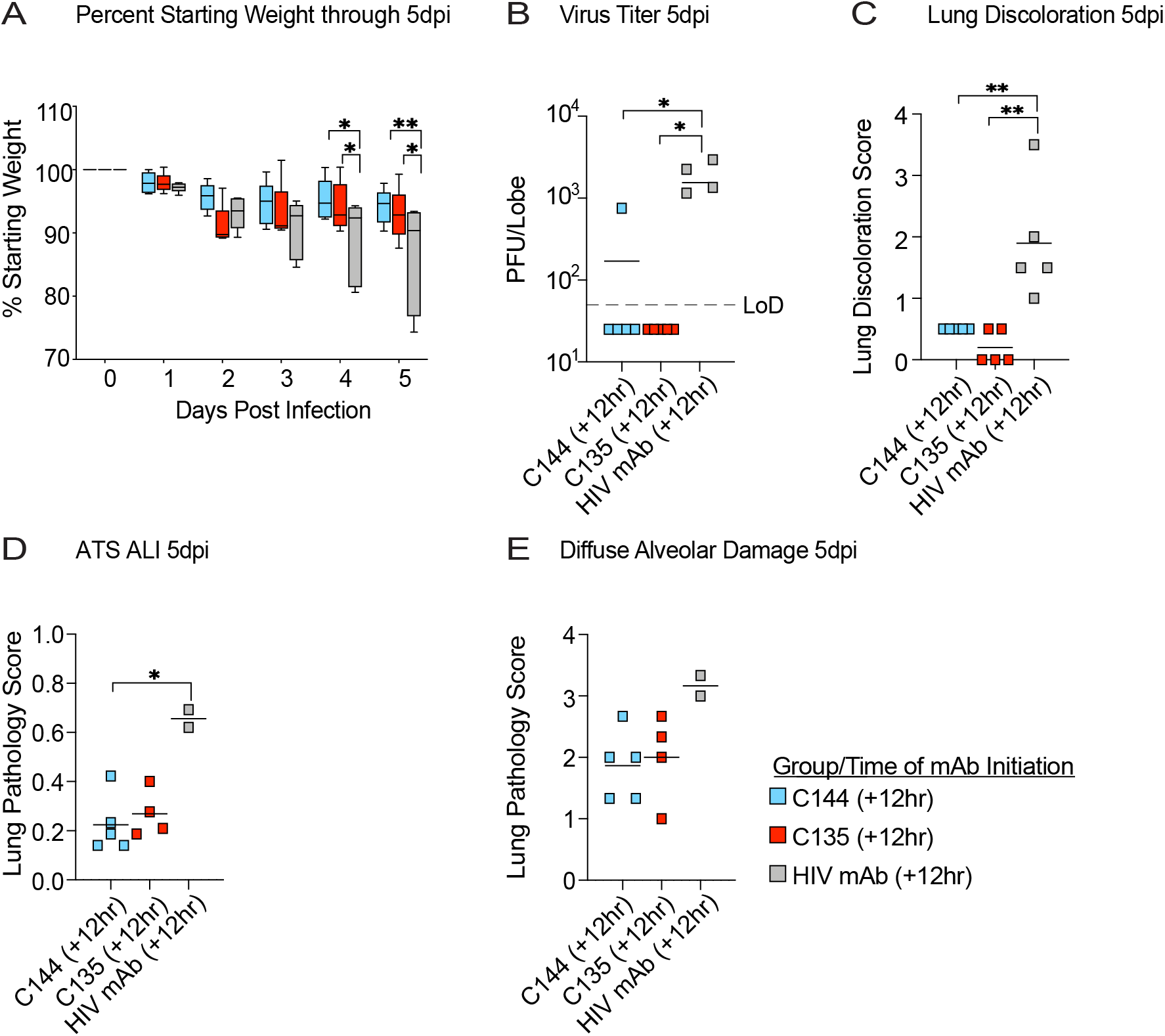
The therapeutic efficacy of mAb monotherapy against SARS-CoV-2 in mice treated at 12 hours post infection. (A) % starting weight in therapeutically treated mice with C144, C135, or an HIV mAb at 12 hours post infection. (B) Lung viral titers in therapeutically treated mice at 12 hours post infection. (C) Lung discoloration score in therapeutically treated mice at 12 hours post infection. (D-E) Lung pathology in therapeutically treated mice at 12 hours post infection. P values are from a 2-way ANOVA after Sidak’s multiple comparisons test.

**Figure S5.**
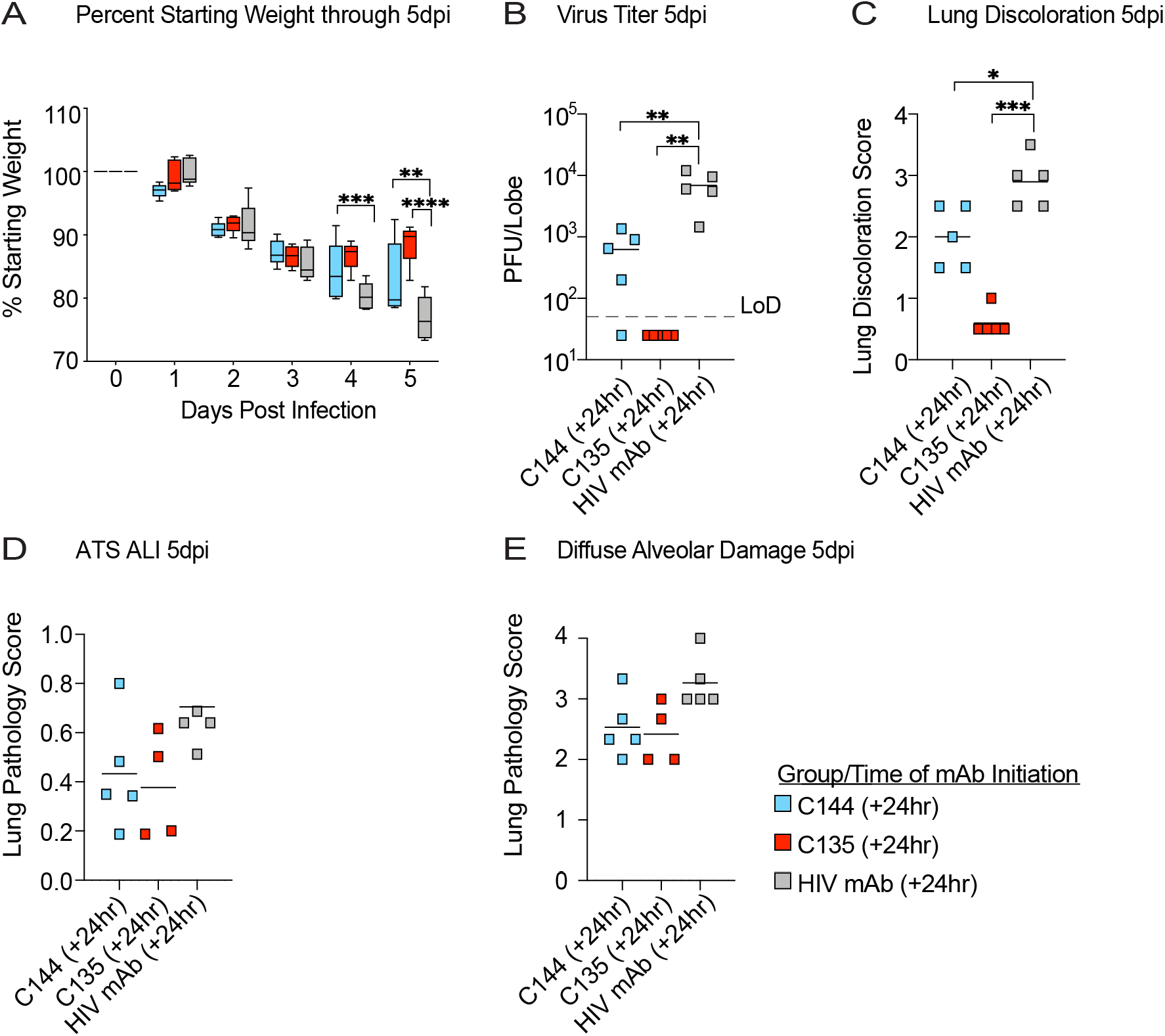
The therapeutic efficacy of mAb monotherapy against SARS-CoV-2 in mice treated at 24 hours post infection. (A) % starting weight in therapeutically treated mice with C144, C135, or an HIV mAb at 24 hours post infection. (B) Lung viral titers in therapeutically treated mice at 24 hours post infection. (C) Lung discoloration score in therapeutically treated mice at 24 hours post infection. (D-E) Lung pathology in therapeutically treated mice at 24 hours post infection. P values are from a 2-way ANOVA after Sidak’s multiple comparisons test.

**Figure S6.**
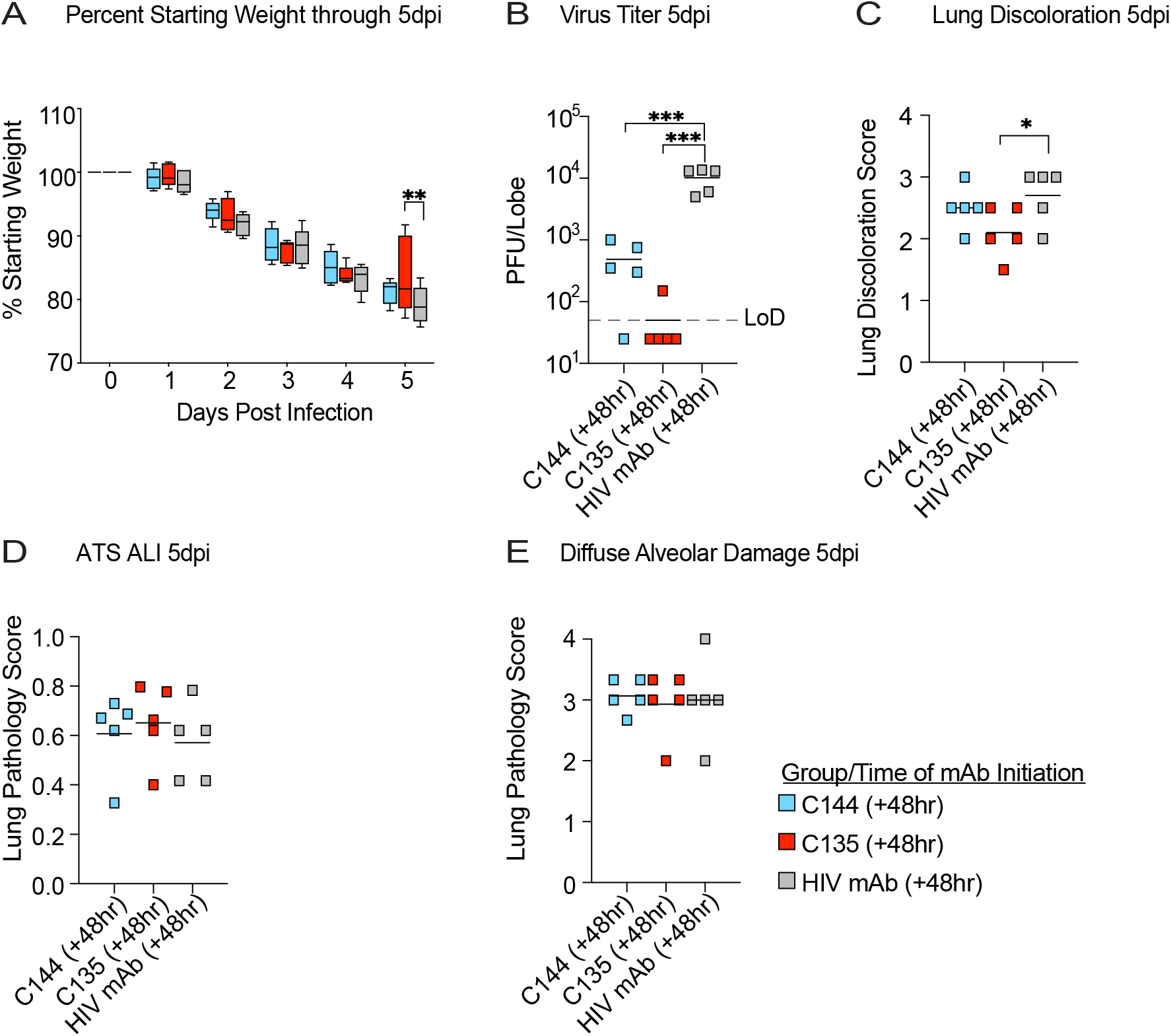
The therapeutic efficacy of mAb monotherapy against SARS-CoV-2 in mice treated at 48 hours post infection. (A) % starting weight in therapeutically treated mice with C144, C135, or an HIV mAb at 48 hours post infection. (B) Lung viral titers in therapeutically treated mice at 48 hours post infection. (C) Lung discoloration score in therapeutically treated mice at 48 hours post infection. (D-E) Lung pathology in therapeutically treated mice at 48 hours post infection. P values are from a 2-way ANOVA after Sidak’s multiple comparisons test.

**Figure S7.**
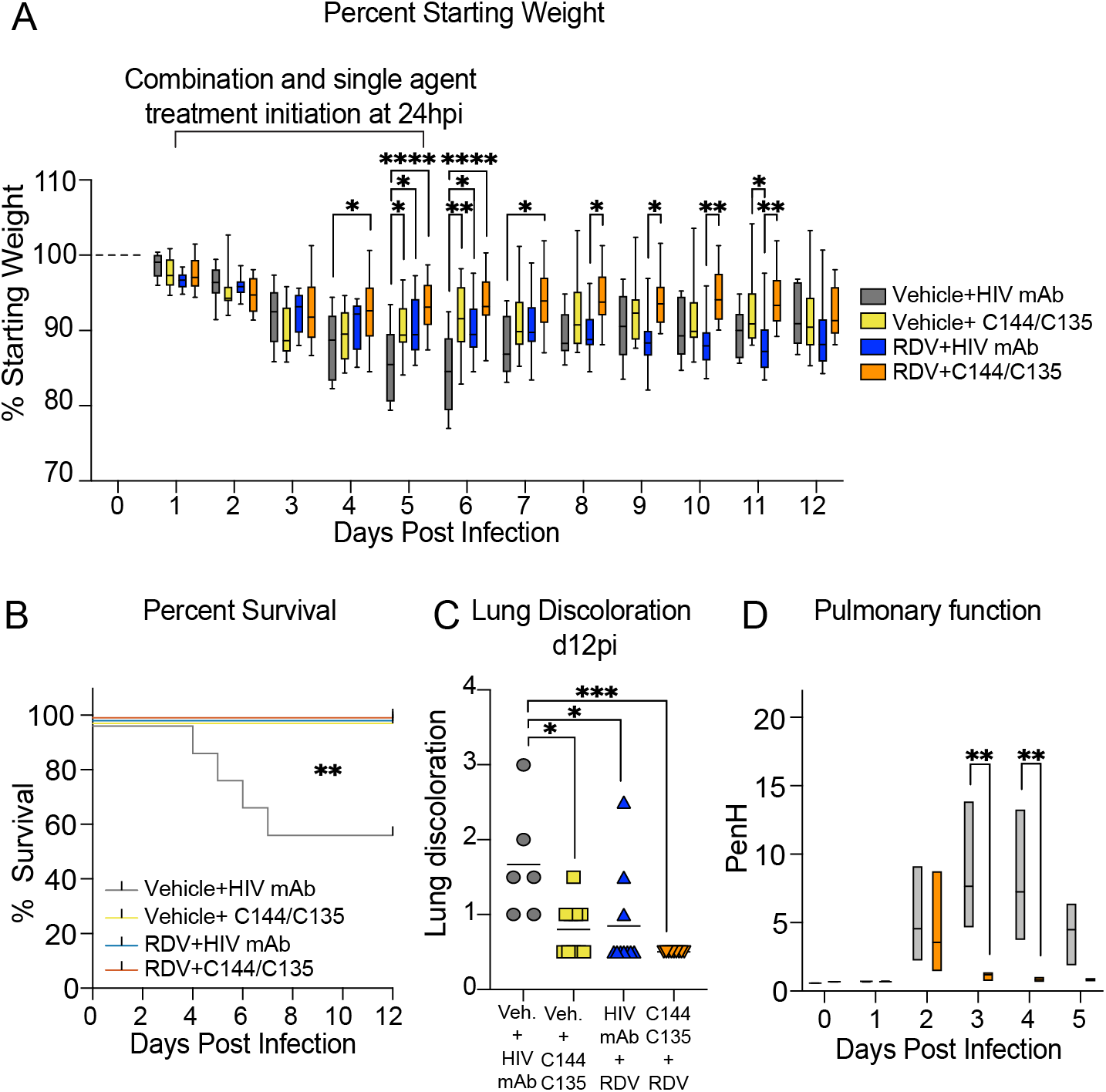
The therapeutic efficacy of RDV and mAbs as single agents and in combination at 24 hours post infection in SARS-CoV-2-infected mice. (A) % starting weight in therapeutically treated mice with vehicle + HIV mAb, vehicle + C144 + C135, RDV + HIV mAb, and RDV + C144 + C135 at 24 hours post infection through day 12. From left to right, grey bars denote vehicle/control mAb treated mice, yellow bars denote vehicle/mAb therapeutic treatment, blue bars denote RDV/control mAb therapeutic treatment, and orange bars denote RDV/mAb therapeutic treatment. (B) % mortality in therapeutically treated mice with single agents and combination therapy. (C) Lung discoloration score in therapeutically treated mice with single agents and combination therapy. “Veh.” signifies vehicle treatment. (D) Pulmonary function in therapeutically treated mice with vehicle + HIV mAb and RDV + C144 + C135. P values are from a 2-way ANOVA after Sidak’s multiple comparisons test.

